# Creation and validation of 3D-printed head molds for stereotaxic injections of neonatal mouse brains

**DOI:** 10.1101/2020.12.22.424033

**Authors:** Ethan Chervonski, Asa A. Brockman, Rohit Khurana, Yuhao Chen, Sabrina Greenberg, Madeleine S. Hay, Yifu Luo, Jason Miller, Deanna Patelis, Sarah K. Whitney, Matthew Walker, Rebecca A. Ihrie

**Affiliations:** Departments of Cell and Developmental Biology, Vanderbilt University; Biomedical Engineering, Vanderbilt University; Neurological Surgery, Vanderbilt, University Medical Center

## Abstract

**Background:** An increasing number of rodent model systems use injection of DNA or viral constructs in the neonatal brain. However, approaches for reliable positioning and stereotaxic injection at this developmental stage are limited, typically relying on handheld positioning or molds that must be re-aligned for use in a given laboratory.

**New method:** A complete protocol and open source software pipeline for generating 3D-printed head molds derived from a CT scan of a neonatal mouse head cast, together with a universal adapter that can be placed on a standard stereotaxic stage.

**Results:** A series of test injections with adenovirus encoding red fluorescent protein were conducted using original clay molds and newly generated 3D printed molds. Several metrics were used to compare spread and localization of targeted injections.

**Comparison with existing methods:** The new method of head mold generation gave comparable results to the field standard, but also allowed the rapid generation of additional copies of each head mold with standardized positioning of the head each time.

**Conclusions:** This 3D printing pipeline can be used to rapidly develop a series of head molds with standardized injection coordinates across multiple laboratories. More broadly, this pipeline can easily be adapted to other perinatal ages or species.

## Introduction

Techniques for *in vivo* genetic manipulation and lineage tracing in neonatal mouse brains have been used with increasing frequency in neural stem cell and brain tumor research (Feliciano et al., 2013; Hoeman et al., 2019; Kim et al., 2019; Merkle et al., 2014; Nagaraja et al., 2017; Qin et al., 2017). These techniques often involve injections of viral vectors, plasmids, small molecules, or cells into specific anatomical regions of the neonatal rodent brain. Injection techniques range in precision from freehand injections to the use of three-dimensional stereotaxic coordinates.

In particular, stereotaxic injection, which is widely used in adult rodent brains, allows researchers to reproducibly target neuroanatomical structures with high spatial precision (Bielefeld et al., 2017; Cetin et al., 2006; Merkle et al., 2007). Brain atlases (Franklin and Paxinos, 2013; Lein et al., 2007) provide useful three-dimensional coordinate systems to locate adult brain structures with respect to a point of reference, such as the bregma, lambda, and interaural line. However, the equivalent methods for neonatal injections require the generation and orientation of head molds, which have the potential for high variability between preparations. Current experimental tools secure neonate heads with customized, laboratory-specific head molds made from malleable clay impressed with casts of neonate heads (Merkle et al., 2007; Merkle et al., 2004). Each new iteration of constructed head molds typically necessitates the re-derivation of successful stereotaxic coordinates due to changes in cast positioning while stamping the mold. This process can be time consuming and can confound the ability to reproduce experiments. Designing head molds that are 3D printed and compatible with existing stereotaxic stages would strengthen the degree of reproducibility across experimental runs and allow more straightforward sharing of injection procedures by collaborating groups.

Here, a 3D-printed apparatus was designed for stereotaxic injection of the neonatal mouse. The design includes a universal stage, which can be secured with ear bars traditionally used on adult mouse or rat injection rigs, and a set of modular, swappable head molds that are customized to specific pup weights and head orientations. This system was validated through a comparative analysis of stereotaxic injections with 3D-printed and clay head molds in the dorsal and ventral mouse ventricular-subventricular zone (V-SVZ), a brain region where targeting with clay head molds has been previously reported. The data presented here provide proof of feasibility for a system that can be adapted to reproducibly inject many other brain regions.

## Materials and Methods

### Construction of 3D-printed head mold

To construct a 3D-printed head mold for a neonate of a given mass, a neonate cast was first developed. When the mouse achieved its desired mass (e.g. CD-1 neonate typically achieves a mass of 1.5-2.1 g at postnatal day 1-2), the animal was euthanized by hypothermia. The neonate was submerged in 3% agarose, and after solidifying, the pup was carefully removed from the agarose to create a mold. The mold was refilled with Hygenic® Repair Resin material and permitted sufficient time to harden. The head of the neonate cast was then imaged with a microCT scan.

The resulting raw scanner file was funneled through the MicroCT EVAL program IPLV6_SEG_CONVERT_STL.COM, streamlining the downsampling and conversion of the raw scanner file to an STL file.

Prior to inverting and re-orienting the scanned head into a mold, the mouse STL file was preprocessed with Meshfix (Attene, 2010) https://github.com/MarcoAttene/MeshFix-V2.1) to remove extraneous holes and lines in the model.

The resulting STL file was placed in a folder with an OpenSCAD script (https://github.com/ihrie-lab/head_mold/blob/master/mold_generator.scad) designed to invert the contour of the head into an imprinted head mold. The default parameters “model_rotate” and “model_translate” were modified to reorient the neonate head as desired on the block. The final head mold designs, which featured two different head mold trays for left and right hemisphere injections, were rendered and saved for printing (https://github.com/ihrie-lab/head_mold/tree/master/head%20mold%20trays). The head molds were printed with polylactic acid (PLA) on Makerbot 3D printers.

Additionally, a universal stage (https://github.com/ihrie-lab/head_mold/blob/master/universal_stage.stl) to secure the swappable head molds to the stereotaxic rig was printed with polylactic acid (PLA) on Makerbot 3D printers.

### Construction of clay head mold

Clay head molds were similarly constructed using neonate casts, following the procedure detailed in Merkle and Tramontin (Merkle et al., 2004). The acrylic neonate cast was laid sideways, and one hemisphere was impressed into a small mound of Sculpey® polymer clay, which was then baked to harden. The other side of the neonate cast was impressed into a different mound of Sculpey® polymer clay, creating separate head molds for each injection hemisphere. Each hardened clay mold was then trimmed and embedded in modeling clay on top of an inverted Petri dish for stability.

### Viral injection of mouse brains with 3D-printed and clay head molds

To compare the head molds, wildtype 1.6 – 1.9 g (age P1-P2) CD-1 mice were injected with human adenovirus type 5 (dE1/E3) expressing mCherry protein under the control of a CMV promoter (Vector Biolabs) using either the 3D-printed or clay head mold. All mice were injected on head molds designed from the cast of a 1.7 g mouse. Injections were conducted on a custom stereotaxic injection apparatus with coordinates previously deemed successful for targeting the dorsal and ventral V-SVZ with clay head molds. Specifically, the injection coordinates targeted radial glial processes in the striatum, thus ultimately labelling cell bodies adjoining the lateral ventricles (Merkle et al., 2004).

Prior to injection, wildtype CD-1 neonates were protected in gauze and a nitrile sleeve and placed on ice for 5-10 minutes to induce anesthesia by hypothermia. The neonate was then laid on its side in a clay or 3D-printed head mold such that one side of the head was securely stabilized in the contours of the mold, leaving the other hemisphere of the head exposed for injection. Transparent tape was used to stretch the skin of the head taut and stabilize the pup in the mold. The clay and 3D-printed head molds were fastened to the stereotaxic stage by tape and ear bars, respectively. A beveled injection needle, pre-loaded with mineral oil, was loaded with the virus, ensuring no air bubbles at the oil/aqueous interface. The needle was attached to the apparatus at a 45° angle above the horizontal plane of the injection stage. The tip of the injection needle was centered on the neonate’s eye, and this position was calibrated as (x = 0 mm, y = 0 mm, z = 0 mm) prior to adjustment to the desired coordinates for dorsal or ventral V-SVZ targeting (Table 1). 100 nl of virus were injected. The procedure was repeated for injection into the other hemisphere. The neonate was rewarmed following injection and returned to its dam.

**Table 1:** Stereotaxic coordinates for subregion-specific targeting of V-SVZ on 1.7 +/− 0.2 g mice

Supplementary Figure and Tables
|  | Stereotaxic Coordinates (from center of eye) |  |  |
| --- | --- | --- | --- |
| Targeting Location | Posterior (mm) | Lateral (mm) | Depth (mm) |
| Dorsal | 2.0 | 1.5 | 2.5 |
| Ventral | 2.0 | 3.0 | 3.6 |

### Post-injection mouse brain fixation and harvesting

Mice were euthanized 3-4 days following injection (postnatal day 4-6) by lethal overdose of 100 μl of 2.5% Avertin. After passing a toe pinch test, they were perfused with 2-3 mL of 0.9% saline solution, followed by 2-3 mL of 4% paraformaldehyde at pH 7.2 containing 0.1M phosphate buffer. The brain was removed from the skull and fixed overnight in 4% paraformaldehyde. Brains were subsequently sunk in 30% sucrose solution at 4°C prior to cryosectioning.

### Post-fix mouse brain processing and fluorescent imaging

Coronal brain sections were sliced serially at 60- or 70-μm thickness from anterior to posterior on a microtome cooling stage regulated by the Physitemp BFS-40MP controller. Sections were mounted directly onto slides in column-major order. Slides were allowed sufficient time to dry and subsequently stored at −20°C with desiccant (Drierite) until immunostaining.

For staining, issue sections were incubated in 200 μl of blocking solution (PBS/1%NDS/1%BSA/0.1% Triton X-100) for 30 minutes at room temperature and subsequently incubated in 200 μl of rabbit anti-RFP antibody (1:1000, MBL PMOO5) overnight at 4°C. The sections were washed and then incubated in 200 μl of donkey anti-rabbit Alexa Fluor 594 antibody (1:1000, Life Technologies A21207) and DAPI (1:10000) for 2-3 hours at room temperature in the dark. Slides were washed, cover slipped with 200 μl of Mowiol mounting media, and stored at −20°C until imaging.

Slides were fluorescently imaged at the Digital Histology Shared Resource of Vanderbilt University Medical Center with the Leica Aperio Versa 200 platform. Images were acquired at 10X magnification with filter sets optimized for DAPI and Texas Red. Sections with localized, small patterns of RFP targeting of V-SVZ regions were imaged at 20X magnification on the LSM 710 META Inverted confocal microscope at the Vanderbilt University Cell Imaging Shared Resource.

### Quantification of spread of injected virus along anterior-posterior axis

The spread of RFP-labelled cells along the anterior-posterior axis of the mouse brain was calculated by a researcher blinded to the type of head mold used. For each hemisphere of the brain, the number of sections containing RFP at the injection site was counted and multiplied by 60 μm or 70 μm, depending on the tissue thickness during sectioning.

### Quantification of percent V-SVZ targeting

The percent of RFP targeting of the V-SVZ was quantified using ImageJ by a researcher blinded to the head mold used. Eligible coronal sections for quantification were limited to V-SVZ-containing sections anterior to the foramen of Monro (position 322 in the Allen Developing Mouse Brain Atlas for a postnatal day 4 pup, https://developingmouse.brain-map.org/experiment/thumbnails/100034998?image_type=nissl). For each eligible section, a pixel-to-micrometer conversion was applied using the scale bar. The coronal tissue section’s entire lateral ventricle was outlined with the freehand line tool, and the ventricle’s perimeter was calculated. A second freehand line tool measurement was used to measure the length along the perimeter that contained RFP-labelled cells in the V-SVZ. The length of targeting was divided by the perimeter of the ventricle and multiplied by 100 to determine the percent V-SVZ targeting for a given section. Percent targeting was averaged among quantified sections of an injected hemisphere.

### Quantification of percent V-SVZ targeting spread along the anterior-posterior axis

For this analysis, brains were anatomically aligned (i.e. designated as the 0-um mark) along the anterior-posterior axis at the first coronal tissue slice in which olfactory areas begin to connect with cortical areas (position 420 in the Allen Developing Mouse Brain Atlas for a postnatal day 4 pup, https://developingmouse.brain-map.org/experiment/thumbnails/100034998?image_type=nissl). The percent V-SVZ targeting of a tissue section was coupled to its posterior distance from this standardized anatomical landmark. Sections without V-SVZ or without targeting of the V-SVZ were assigned values of zero percent V-SVZ targeting for the purpose of this analysis. The spread and location of V-SVZ targeting along the anterior-posterior axis were compared graphically.

## Results

3D-printed head molds were generated using a script-based workflow that inverts and orients microCT scans of mouse head casts into blocks, thus virtually “stamping” head molds (Fig. 1A; https://github.com/ihrie-lab/head_mold/blob/master/mold_generator.scad). Each head mold standardized a head orientation for injection of mice of a given mass (https://github.com/ihrie-lab/head_mold/tree/master/head%20mold%20trays). A 3D-printed universal stage, which was fastened to the ear bars of the stereotaxic rig, was also designed to secure these swappable head molds in a consistent alignment (stage design shown in Fig. 1A; https://github.com/ihrie-lab/head_mold/blob/master/universal_stage.stl). The modular design of the individual 3D-printed head mold trays and the universal stage permits easy exchange and consistent orientation of head molds for different mouse sizes and brain hemispheres of injection. Notably, the universal stage design also contained a cooling compartment for inclusion of a solid coolant pack to maintain lower body temperatures during injection.

**Fig. 1:**
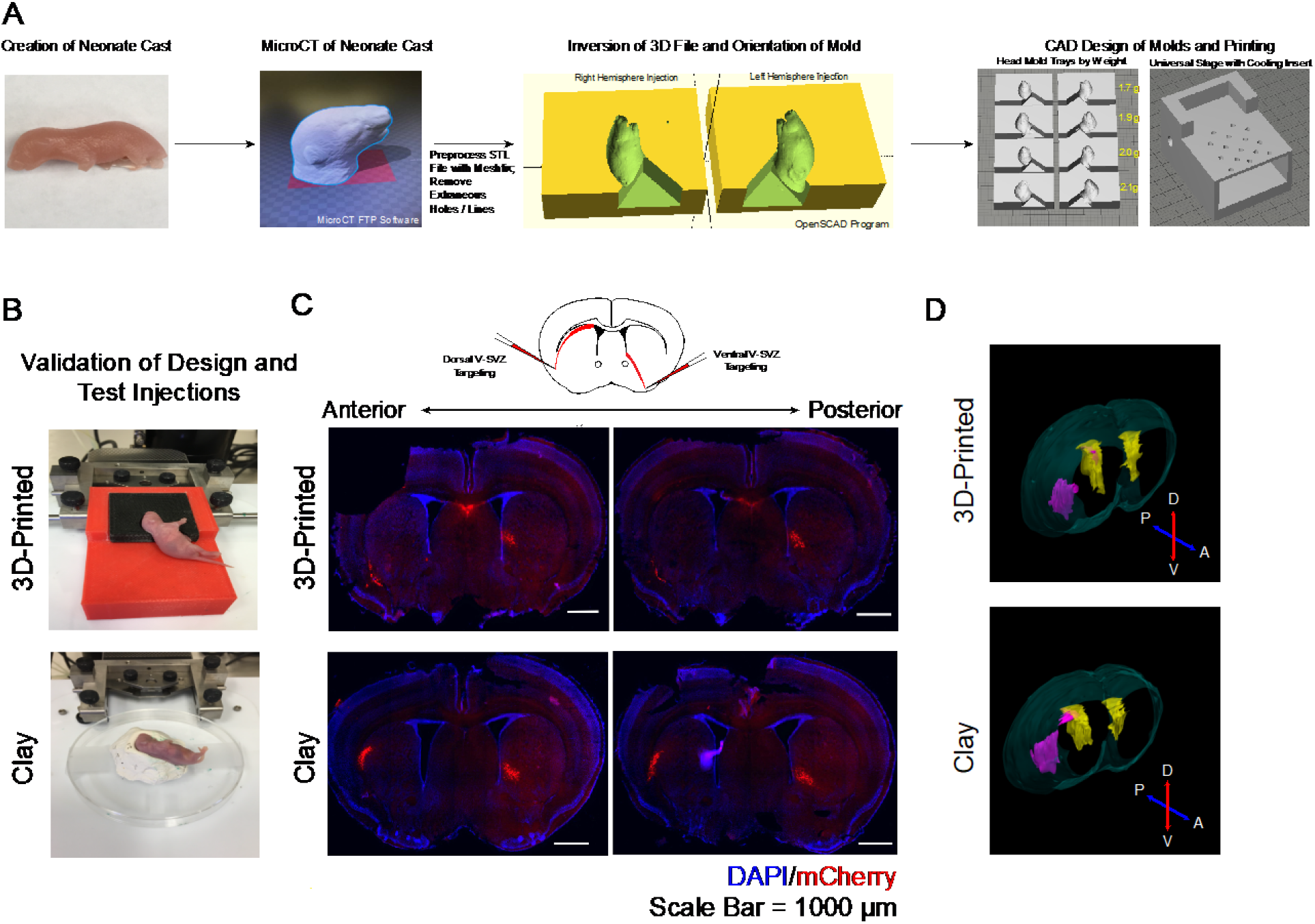
A comparison of injections visualized through fluorescent microscopy and 3D brain reconstructions showed no injection differences. A) 3D-printed head mold construction workflow; B) Representative images of injection set-up with 3D-printed and clay head molds; C) Representative wide field images of serial coronal mouse brain sections (60-μm thickness) with viral targeting of ventral and dorsal V-SVZ through striatal injections, scale bars = 1000 μm; D) Representative 3D reconstructions of mouse brains with dorsal V-SVZ targeting, color scheme: blue = brain surface, yellow = V-SVZ, pink = dorsal injection labelling.

To validate the 3D-printed head mold design, injections targeting the V-SVZ, a neurogenic microenvironment that lines the lateral ventricles (Codega et al., 2014; Ihrie and Alvarez-Buylla, 2011; Morshead and van der Kooy, 2004), with recombinant mCherry adenovirus were scored and compared between the 3D-printed and clay head molds (Fig. 1B). The ability to access one V-SVZ cell subpopulation while leaving other subregions unperturbed, whether through stereotaxic injection, microdissection, or transgenic modification in mice, has allowed researchers to associate differential transcription factor expression, signaling activity, and lineage commitment with cellular positional identity in the V-SVZ (Llorens-Bobadilla et al., 2015; Merkle et al., 2014; Merkle et al., 2007; Mizrak et al., 2019; Rushing and Ihrie, 2016; Rushing et al., 2019; Young et al., 2007). As localized, consistent targeting of the V-SVZ could reveal more refined characterizations of neural stem cell positional identity, injection metrics were designed to measure the spread and reproducibility of targeting dorsal or ventral V-SVZ cell clusters.

Neonatal mice were injected at 1.6-1.9 g body mass (postnatal day 1-2) using striatal coordinates derived for ventral and dorsal V-SVZ targeting. Injection of radial glial processes in the mouse striatum labeled radial glial cell bodies housed in the periventricular region, as expected (Fig. 1 C-D)(Merkle et al., 2004). 26 mice were injected bilaterally for a total of 52 injections. One hemisphere was excluded from all subsequent analysis because the mislocalized injection was too posterior, and one other hemisphere was excluded from analyses requiring V-SVZ quantifications because of section tearing. From the remaining 51 injections, 76.5% showed a detectable RFP-labelled injection site in the striatum. Among the 39 labelled hemispheres studied, a greater percentage of injections with the 3D-printed head mold showed successful targeting of the dorsal V-SVZ—that is, RFP-labelled cell bodies were detectable in the dorsal V-SVZ. Fewer injections with the 3D-printed head mold consistently targeted the ventral V-SVZ, although sufficient targeted hemispheres were available for analysis (Table 2).

**Table 2:**
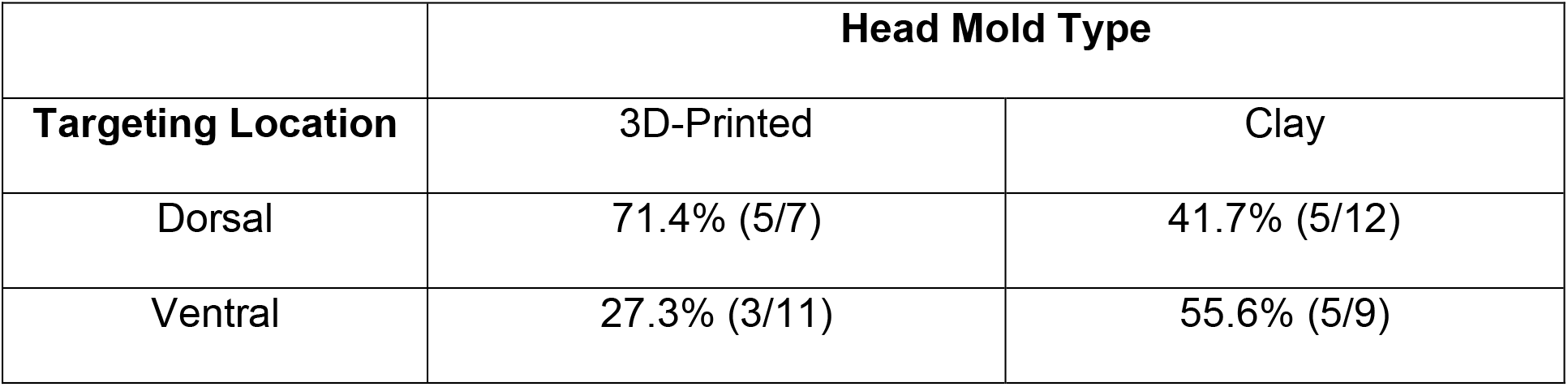
Sμmmary table of percentage of RFP-labelling injections that successfully targeted the V-SVZ (with fraction of successfully targeted brains shown)

The anterior-to-posterior spread of the virus was quantified to compare consistency at the injection site, a feature that would likely not change between head molds. A Wilcoxon rank-sum test comparing injections with 3D-printed and clay head molds revealed no difference in the spread of virus along the anterior-posterior axis for either dorsal (p = 0.0906) or ventral injections (p = 0.1681) (Fig. 2A). Additionally, the percentage of the V-SVZ labelled by the virus and its location/spread along the anterior-posterior axis were also compared as measures for the degree of targeting localization, which is vital to tracking precise groups of neural stem cells. A Wilcoxon rank-sum test comparing percent V-SVZ targeting among successfully targeted injections showed no difference between 3D-printed and clay head molds for dorsal (p = 0.2851) or ventral injections (p = 0.5000) (Fig. 2B). Finally, comparing the spread and location of V-SVZ targeting along the anterior-posterior axis graphically showed that 3D-printed and clay head molds were similar (Fig. 2C-D). Quantitative comparisons of brain injections with Ad-mCherry virus on 1.7 +/− 0.2 g mice therefore revealed that injections with the 3D-printed head mold were as precise and consistent as those with the clay head mold and, in the dorsal V-SVZ, yielded a higher number of successful injection events.

**Fig. 2:**
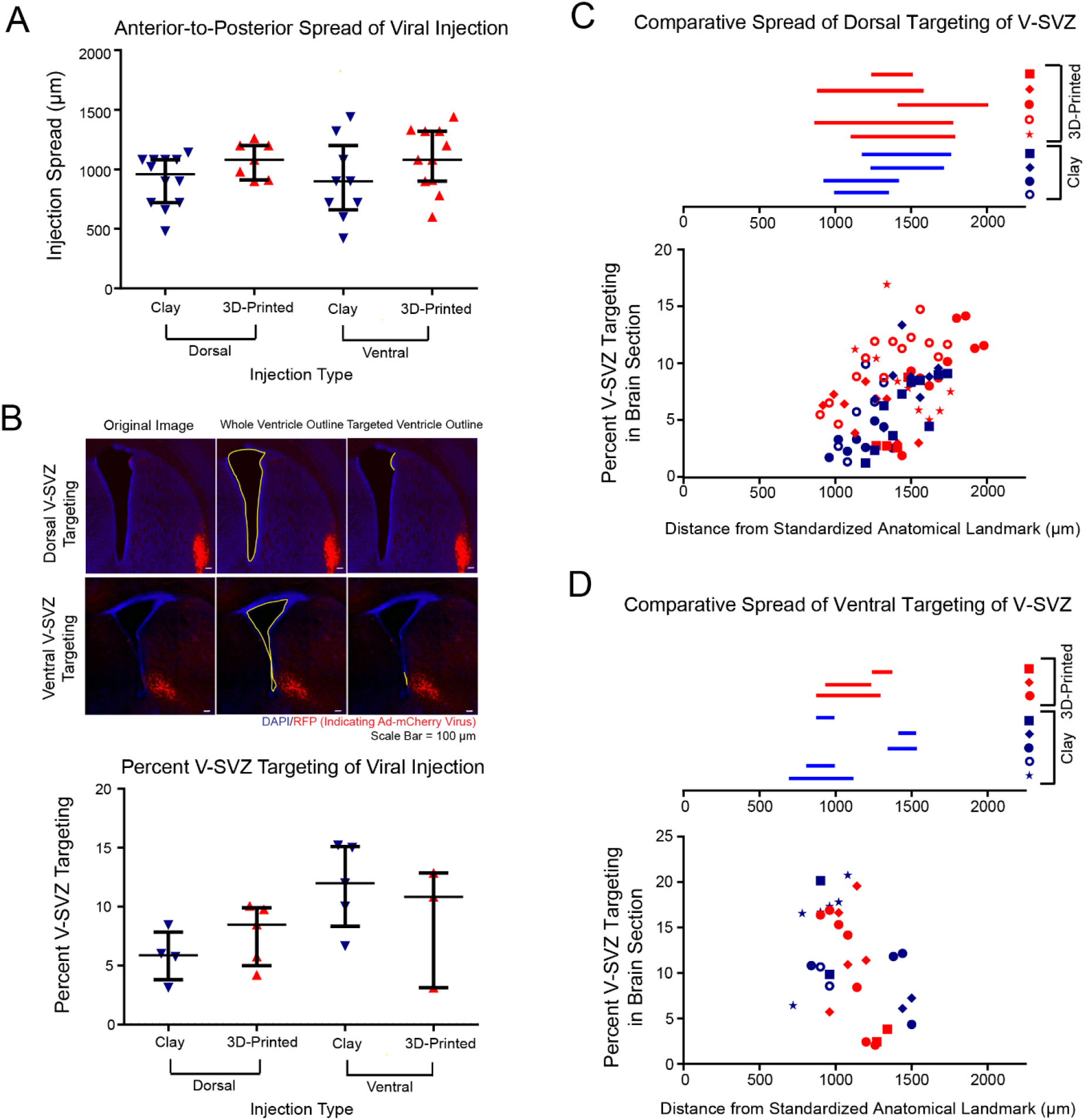
A quantitative comparison of viral injection features revealed no differences between injections with the 3D-printed and clay head molds on 1.6-1.9 g mice. A) Data plot comparing anterior-to-posterior spread of dorsal and ventral injections with 3D-printed and clay head molds, median and interquartile range shown; B) Top: schematic of image analysis technique for percent V-SVZ computation (targeted ventricle outline/whole ventricle perimeter x 100), scale bars = 100 μm; Bottom: Data plot comparing percent V-SVZ targeting of dorsal and ventral injections with 3D-printed and clay head molds, median and interquartile range shown; C) Distribution of dorsal V-SVZ targeting from a standardized anatomical point (represented as 0 μm) along the anterior-posterior axis, each shape indicates a different brain hemisphere of injection; D) Distribution of ventral V-SVZ targeting from a standardized anatomical point (represented as 0 μm) along the anterior-posterior axis, each shape indicates a different brain hemisphere of injection

## Discussion

Stereotaxic injection allows researchers to access highly specific neuroanatomical locations for genetic perturbation, drug administration, or lesion generation. However, such injections can be time-consuming and follow up timepoints can be lengthy, meaning that reproducibility in targeting is highly desirable in reducing the number of unsuccessful or mistargeted replicates. In the case of this study, stereotaxic injection permitted testing of two head mold types when targeting cell subpopulations of the neonatal mouse V-SVZ.

A standardized workflow for head molds permits the maintenance of stereotaxic targeting coordinates for a particular anatomical landmark without re-derivation when head molds are newly produced. The construction of clay head molds is susceptible to problems in consistent alignment and orientation of the mouse cast while impressing it into the clay mold, requiring the re-derivation of stereotaxic coordinates for each newly produced head mold. Additionally, the 3D-printed universal stage included in this package consistently aligns and orients the head molds relative to the stereotaxic rig, whereas the current clay head mold cannot be securely fastened to the stage and therefore may change with each placement.

The results of comparative image analyses of viral labelling in the V-SVZ for injections using the two head mold construction workflows indicate that injections with 3D-printed head molds are comparable to injections with clay head molds. The anterior-posterior spread of virus, percent V-SVZ targeting, and spread/location of V-SVZ targeting along the anterior-posterior axis were not significantly different between head molds across injection locations. Similar results have been noted on a subset of 2.1 +/− 0.2 g mice using head molds and stereotaxic coordinates tailored towards a 2.1 g mouse (Supplementary Fig. 1). While the sample size for this subset of mice was smaller, no difference in targeting success metrics was seen between head molds.

The observed variations in injection metrics for both clay and 3D-printed head molds may be due to multiple sources. First, the derivation of stereotaxic coordinates was conducted on clay head molds. Therefore, the coordinates, especially for ventral injections targeting a highly specific subregion, were likely not optimally suited for the 3D-printed head orientation. Additionally, neonates’ mass varied from 1.6 to 1.9 g, but the same 1.7 g-mouse head mold/coordinates were used for all injections. V-SVZ subregion coordinates may vary across this small mass range.

3D-printed head molds provide additional value in their ability to standardize stereotaxic coordinates for a given anatomical landmark across newly produced head molds. While stereotaxic atlases exist to provide clear locations of adult anatomical structures for quick reference, injecting neonates depends on a researcher’s ability to orient the head carefully relative to the stereotaxic rig, a nontrivial task given the small size of neonates. The publication of 3D-printed head mold CAD files for wider use is a step towards a practical standard in the field for precise neonatal injections of any anatomical landmark. This would reduce the requirement for arduous customization of head molds and re-derivation of coordinates for rodent brain structures as new projects arise.

## Acknowledgements

The authors thank the Ihrie and Irish labs for helpful discussions on data and figure organization and Dr. Jose Maldonado and Dr. Shawn Sorrells for help with 3D brain reconstructions using Neurolucida® software. Whole slide imaging of mouse brain tissue was performed at the Digital Histology Shared Resource at Vanderbilt University Medical Center (www.mc.vanderbilt.edu/dhsr). Confocal microscopy of mouse brain tissue was performed through the use of the Vanderbilt University Cell Imaging Shared Resource (supported by NIH grants S10 1S10OD021630-01, CA68485, DK20593, DK58404, DK59637, and EY08126). This work was supported by the Littlejohn Fellowship/Vanderbilt Undergraduate Summer Research Program (E.C.), R01 NS096238 (R.A.I), DOD W81XWH-16-1-0171 (R.A.I.), and the Vanderbilt School of Engineering (M.W.).

## Author Contributions

E. Chervonski: conceptualization, methodology, validation, investigation, formal analysis, visualization, writing – original draft, review and editing

A.A. Brockman: conceptualization, methodology, investigation, validation, writing – review and editing

R. Khurana: validation, data curation

Y. Chen: methodology, software

S. Greenberg: methodology, software

M.S. Hay: methodology, software

Y. Luo: methodology, software

J. Miller: methodology, software

D. Patelis: methodology, software

S.K. Whitney: methodology, software

M. Walker III: supervision, funding acquisition

R.A. Ihrie: conceptualization, supervision, methodology, funding acquisition, visualization, writing – review and editing

## Supplementary Figure and Tables

**Supplementary Fig. 1:**
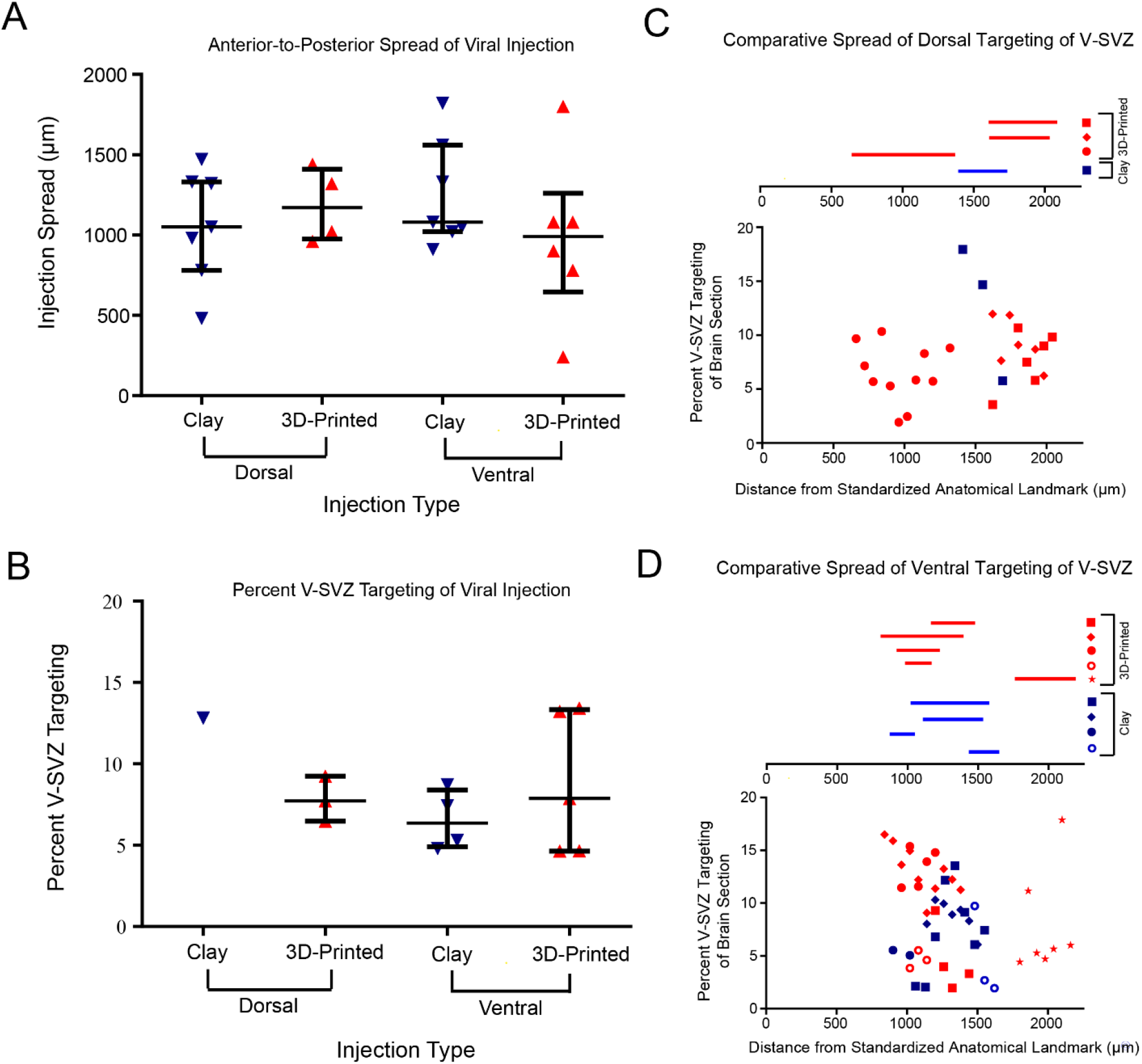
A quantitative comparison of viral injection features revealed no differences between injections with the 3D-printed and clay head molds on 2.0-2.3 g mice. A) Data plot comparing anterior-to-posterior spread of dorsal and ventral injections with 3D-printed and clay head molds, median and interquartile range shown; B) Data plot comparing percent V-SVZ targeting of dorsal and ventral injections with 3D-printed and clay head molds, median and interquartile range shown; C) Distribution of dorsal V-SVZ targeting from a standardized anatomical point (represented as 0 μm) along the anterior-posterior axis, each shape indicates a different brain hemisphere of injection; D) Distribution of ventral V-SVZ targeting from a standardized anatomical point (represented as 0 μm) along the anterior-posterior axis, each shape indicates a different brain hemisphere of injection.

## References

Allen Developing Mouse Brain Atlas, available from:https://developingmouse.brain-map.org/

Attene, M. (2010). A lightweight approach to repairing digitized polygon meshes. The Visual Computer 26, 1393–1406.

Bielefeld, P., Sierra, A., Encinas, J.M., Maletic-Savatic, M., Anderson, A., and Fitzsimons, C.P. (2017). A Standardized Protocol for Stereotaxic Intrahippocampal Administration of Kainic Acid Combined with Electroencephalographic Seizure Monitoring in Mice. Frontiers in neuroscience 11, 160.

Cetin, A., Komai, S., Eliava, M., Seeburg, P.H., and Osten, P. (2006). Stereotaxic gene delivery in the rodent brain. Nat Protoc 1, 3166–3173.

Codega, P., Silva-Vargas, V., Paul, A., Maldonado-Soto, A.R., Deleo, A.M., Pastrana, E., and Doetsch, F. (2014). Prospective identification and purification of quiescent adult neural stem cells from their in vivo niche. Neuron 62, 545–559.

Feliciano, D.M., Lafourcade, C.A., and Bordey, A. (2013). Neonatal subventricular zone electroporation. J Vis Exp.

Franklin, K.B.J., and Paxinos, G. (2013). Paxinos and Franklin’s The mouse brain in stereotaxic coordinates, Fourth edition. edn (Amsterdam: Academic Press, an imprint of Elsevier).

Hoeman, C.M., Cordero, F.J., Hu, G., Misuraca, K., Romero, M.M., Cardona, H.J., Nazarian, J., Hashizume, R., McLendon, R., Yu, P., et al. (2019). ACVR1 R206H cooperates with H3.1K27M in promoting diffuse intrinsic pontine glioma pathogenesis. Nat Commun 10, 1023.

Ihrie, R.A., and Alvarez-Buylla, A. (2011). Lake-front property: a unique germinal niche by the lateral ventricles of the adult brain. Neuron 70, 674–686.

Kim, G.B., Rincon Fernandez Pacheco, D., Saxon, D., Yang, A., Sabet, S., Dutra-Clarke, M., Levy, R., Watkins, A., Park, H., Abbasi Akhtar, A., et al. (2019). Rapid Generation of Somatic Mouse Mosaics with Locus-Specific, Stably Integrated Transgenic Elements. Cell 179, 251–267 e224.

Lein, E.S., Hawrylycz, M.J., Ao, N., Ayres, M., Bensinger, A., Bernard, A., Boe, A.F., Boguski, M.S., Brockway, K.S., Byrnes, E.J., et al. (2007). Genome-wide atlas of gene expression in the adult mouse brain. Nature 445, 168–176.

Llorens-Bobadilla, E., Zhao, S., Baser, A., Saiz-Castro, G., Zwadlo, K., and Martin-Villalba, A. (2015). Single-Cell Transcriptomics Reveals a Population of Dormant Neural Stem Cells that Become Activated upon Brain Injury. Cell Stem Cell 17, 329–340.

Merkle, F.T., Fuentealba, L.C., Sanders, T.A., Magno, L., Kessaris, N., and Alvarez-Buylla, A. (2014). Adult neural stem cells in distinct microdomains generate previously unknown interneuron types. Nat Neurosci 17, 207–214.

Merkle, F.T., Mirzadeh, Z., and Alvarez-Buylla, A. (2007). Mosaic organization of neural stem cells in the adult brain. Science 317, 381–384.

Merkle, F.T., Tramontin, A.D., Garcia-Verdugo, J.M., and Alvarez-Buylla, A. (2004). Radial glia give rise to adult neural stem cells in the subventricular zone. Proc Natl Acad Sci U S A 101, 17528–17532.

Mizrak, D., Levitin, H.M., Delgado, A.C., Crotet, V., Yuan, J., Chaker, Z., Silva-Vargas, V., Sims, P.A., and Doetsch, F. (2019). Single-Cell Analysis of Regional Differences in Adult V-SVZ Neural Stem Cell Lineages. Cell Rep 26, 394–406 e395.

Morshead, C.M., and van der Kooy, D. (2004). Disguising adult neural stem cells. Curr Opin Neurobiol 14, 125–131.

Nagaraja, S., Vitanza, N.A., Woo, P.J., Taylor, K.R., Liu, F., Zhang, L., Li, M., Meng, W., Ponnuswami, A., Sun, W., et al. (2017). Transcriptional Dependencies in Diffuse Intrinsic Pontine Glioma. Cancer Cell 31, 635–l652 e636.

Qin, E.Y., Cooper, D.D., Abbott, K.L., Lennon, J., Nagaraja, S., Mackay, A., Jones, C., Vogel, H., Jackson, P.K., and Monje, M. (2017). Neural Precursor-Derived Pleiotrophin Mediates Subventricular Zone Invasion by Glioma. Cell 170, 845–859 e819.

Rushing, G., and Ihrie, R.A. (2016). Neural stem cell heterogeneity through time and space in the ventricular-subventricular zone. Front Biol (Beijing) 11, 261–284.

Rushing, G.V., Brockman, A.A., Bollig, M.K., Leelatian, N., Mobley, B.C., Irish, J.M., Ess, K.C., Fu, C., and Ihrie, R.A. (2019). Location-dependent maintenance of intrinsic susceptibility to mTORC1-driven tumorigenesis. Life Sci Alliance 2.

Young, K.M., Fogarty, M., Kessaris, N., and Richardson, W.D. (2007). Subventricular zone stem cells are heterogeneous with respect to their embryonic origins and neurogenic fates in the adult olfactory bulb. J Neurosci 27, 8286–8296.

